# Association of Model Neurotransmitters with Lipid Bilayer Membranes

**DOI:** 10.1101/822189

**Authors:** B. Josey, F. Heinrich, V. Silin, M. Lösche

## Abstract

Aimed to reproduce the results of electrophysiological studies of synaptic signal transduction, conventional models of neurotransmission are based on the specific binding of neurotransmitters to ligand-gated receptor ion channels. However, the complex kinetic behavior observed in synaptic transmission cannot be reproduced in a standard kinetic model without the ad hoc postulation of additional conformational channel states. On the other hand, if one invokes unspecific neurotransmitter adsorption to the bilayer—a process not considered in the established models—the electrophysiological data can be rationalized with only the standard set of three conformational receptor states that also depend on this indirect coupling of neurotransmitters via their membrane interaction. Experimental verification has been difficult because binding affinities of neurotransmitters to the lipid bilayer are low. We quantify this interaction with surface plasmon resonance to measure equilibrium dissociation constants in neurotransmitter membrane association. Neutron reflectometry on artificial membranes reveals the structural aspects of neurotransmitters association with zwitterionic and anionic bilayers. We establish that serotonin interacts non-specifically with the membrane at physiologically relevant concentrations whilst GABA (γ-aminobutyric acid) does not. Surface plasmon resonance shows that serotonin adsorbs with millimolar affinity and neutron reflectometry shows that it penetrates the membrane deeply whereas GABA is excluded from the bilayer.

**Significance:** Receptor ion channels in the postsynaptic membrane and their neurotransmitter agonists enable fast communication between neuronal cells. Electrophysiology studies reveal surprisingly complex kinetics that apparently require a variety of protein conformational states for their quantitative interpretation, but an alternate hypothesis invoking neurotransmitter membrane association reduces the complexity of the underlying reaction schemes significantly. While their affinity may be low, and is hard to quantify experimentally, neurotransmitter membrane association can be relevant because of their large temporary concentration in the synaptic cleft. With thermodynamic and structural measurements we quantify membrane-bound states of serotonin, establishing this neurotransmitter as membrane-affine, whereas the affinity of the more hydrophilic GABA is too low to register in our sensitivity-optimized measurement techniques.

## Introduction

Synaptic transmission between neurons is mediated by ligand-gated ion channels (LGIC) embedded in the postsynaptic membrane. These proteins selectively allow either cations (*e.g.* acetylcholine and glutamate receptors) or anions (*e.g.* γ-aminobutyric acid (GABA) and glycine receptors) to flow into the postsynaptic neuron (1). Conventional kinetic reaction schemes require a surprisingly large number of protein conformational states (2–6), yet do not accurately model experimental results (7). For example, desensitization of their ion-conducting conformation decreases with complex kinetics, requiring multiple decay exponentials for data modeling. Although the molecular origin of this complex response is not known, it must be essential to the proper functioning of the central nervous system, since organisms with mutations that alter desensitization kinetics exhibit major phenotypic changes detrimental to the health of the organism (1). Addressing these issues, an alternative hypothesis has been developed which simplifies the kinetic models that describe this complex gating behavior. It posits that neurotransmitters which activate postsynaptic receptors through binding to specific sites also modulate receptor activity indirectly (8–10). This hypothesis invokes NT interactions with the postsynaptic lipid bilayer surrounding the receptor proteins, which may alter its physical properties and in turn modulate receptor activity (11). For a specific case, the interaction of GABA with its receptor GABA_A_, a minimal kinetic scheme based on these ideas described experimental agonist-induced and anesthetic-induced activation-deactivation traces of ion flow across the membrane remarkably well (9). Additional support for this hypothesis came from the observation that the response of a receptor to its cognate NT is altered by the presence of a *different*, unrelated NT (12). While this provided an impressive validation of the unconventional receptor activation concept, it does not directly demonstrate that NT-membrane interactions do indeed occur.

Various experimental approaches to measure non-specific NT binding to lipid bilayers have been reported. Such studies showed that the cationic acetylcholine and the zwitterionic GABA and glycine neurotransmitters bind to anionic membranes containing phosphatidylserine (PS) or phosphatidylglycerol (PG) (13) and that the aromatic 5-hydroxytryptamine (5-HT; serotonin) has a greater partitioning coefficient into such membranes than polar neurotransmitters (14). Atomistic molecular dynamics (MD) simulations suggested that serotonin adsorbs to the anionic phosphate in phosphatidylcholine (PC) through interaction of its cationic primary amine and is then anchored by aligning its aromatic ring with the hydrocarbon tails. This association has been observed to shield against oxidation in erythrocytes (15). The serotonin derived hormone melatonin associates with planar bilayers in a similar orientation, however this has only been measured qualitatively but not quantitatively (16, 17). Another study indicated that the electrostatic interaction of dopamine with the inner leaflet of presynaptic vesicles must be screened to prevent NT aggregation and thus facilitate synaptic release (18).

In MD simulations, hydrophilic NTs (*e.g.* GABA and serine) are generally observed to remain in the solution adjacent to a membrane or interact weakly with charged headgroups, while hydrophobic species (*e.g.* serotonin and dopamine) partition into the bilayer and often localize near the lipid phosphates (19–21). Postila and coworkers studied a broad range of NTs for their membrane interactions and reported a strong correlation between their membrane binding prevalence and the location of the binding sites on their cognate receptors (19). NTs with extracellular binding sites on the receptor showed low affinity; but NTs whose binding sites are hidden within the membrane—typically in the lipid headgroup region—consistently showed high membrane affinities and were classified as membrane-affine. If these observations from simulation can be confirmed with experimental data, this will provide strong support for the hypothesis that the bilayer plays a critical role in the regulation of neurotransmission: while hydrophilic NTs bind directly to their binding sites after diffusing across the synaptic cleft, hydrophobic NTs may first adsorb to the bilayer and subsequently encounter their receptors through two-dimensional diffusion.

Here, we investigate the interactions of two prototypical NTs—serotonin, which was identified as a membrane-affine NT, and GABA, for which simulation results showed low membrane affinity—with model membranes as a function of lipid headgroup charge density. Surface plasmon resonance (SPR) provides quantitative measurements of membrane affinities and shows striking differences between serotonin and GABA. These thermodynamic results are quantitatively consistent with neutron reflection (NR) results by which we study the impact of NTs on membrane structure and localize NTs in the bilayer. In combination, these results show that the aromatic NT serotonin, a cation at physiological pH, binds neutral and charged membranes with millimolar affinity. It intercalates into the bilayer with a distribution that has its highest density at the depth of the phospholipid headgroups, at a position where it can bind its effector protein efficiently. Beyond this specific interaction, it may affect other membrane-embedded proteins, conceivably by modification of the pressure profile across the bilayer. GABA interaction with bilayers was much weaker: the sensitivity of both methods was insufficient for a quantification of surface adsorbed amounts of this NT. However, our SPR measurements established an upper limit of its dissociation constant from the membrane which is above the typical concentration of NTs in the synaptic cleft.

## Materials and Methods

### Materials

1-palmitoyl-2-oleoyl-*sn*-glycero-3-phosphocholine (POPC) and 1-palmitoyl-2-oleoyl-*sn*-glycero-3-phospho-(1-*rac*-glycerol) (POPG, sodium salt) from Avanti Polar Lipids (Alabaster, AL) were used as received. The tether lipid HC18 [Z20-(Z-octadec-9-enyloxy)-3,6,9,12,15,18, 22-heptaoxatetracont-31-ene-1-thiolacetate] (22) was provided by David Vanderah (IBBR, Rockville, MD). Other chemicals, including β-mercaptoethanol (βME), salts and the NTs GABA and serotonin, were from Sigma-Aldrich (St. Louis, MO).

### Preparation of Sparsely-Tethered Bilayer Lipid Membranes (stBLMs)

stBLMs (Fig. 1) were prepared as follows. Self-assembled monolayers (SAMs) composed of βME and HC18 (7:3 in solution) were formed on gold-covered solid substrates, followed by bilayer completion through rapid solvent exchange (23) or vesicle fusion (24). For SPR, we used microscopy slides made of glass (Thermo Fisher Scientific, Waltham, MA) or sapphire (Rubicon, Bensonville, IL). Reflectivity measurements were performed on 3’’ diameter, 5 mm thick silicon waters (El-Cat Inc. Ridgefield Park, NJ). The bare substrates were cleaned in 5 vol% Hellmanex solution (Hellma Analytics, Müllheim, Germany), incubated in sulphuric acid with Nochromix (Godax Laboratories, Cabin John, MD) for 15 minutes, rinsed with ample volumes of ultrapure water (EMD Millipore, Billerica, MA) and ethanol, and dried in an N_2_ stream. They were then coated in a magnetron (Denton Vacuum Discovery 550) with a ≈ 40 Å thick chromium bonding layer, followed by a terminal gold layer that was ≈ 450 Å thick for SPR and ≈ 150 Å thick for NR measurements. In all cases, the RMS interfacial roughness of the gold surfaces were σ < 5 Å.

**Figure 1:**
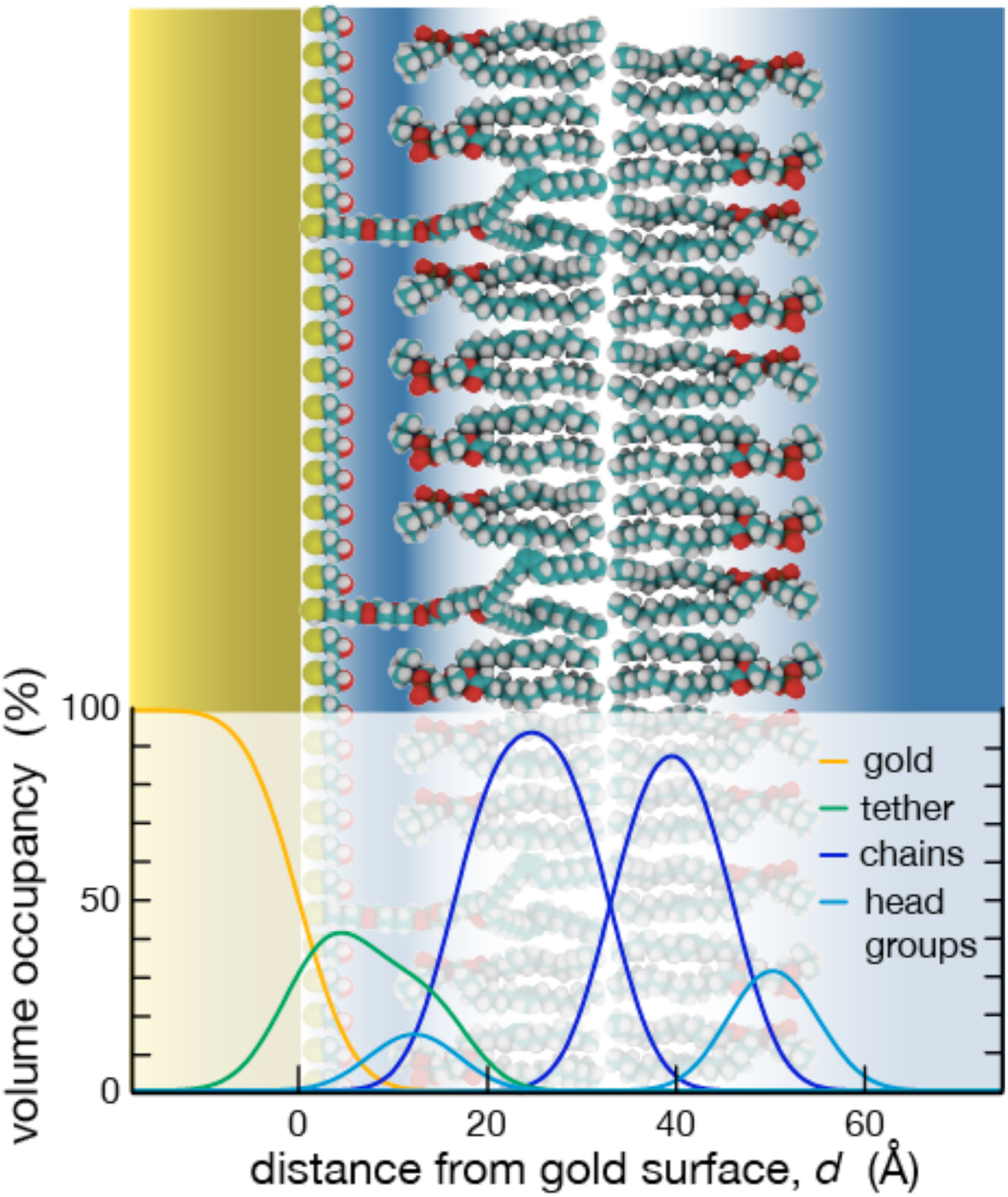
The solid-supported membrane model system used in this study and its structural characterization. A sparsely-tethered bilayer lipid membrane (stBLM) consists of a substrate-proximal lipid layer that is tethered to a gold surface (25, 63) and a distal layer that is fully fluid (64). Such systems have been extensively characterized with NR, and their out-of-plane structure is described in terms of distributions in which the molecular components occupy volume within the the plane of the bilayer, as exemplified in the lower section, and referred to as component volume occupancy, CVO. The proximal lipid layer contains more chain volume than the distal layer because the tether lipids are ether-based and pack more densely than the intervening phospholipids. In the quantitative model, water (distribution not shown) fills any space where the combined CVOs of the organic molecules is less than unity (34, 38).

For bilayer formation on the SAMs through vesicle fusion, used in the NR experiments, stock lipids dissolved in organic solvents were mixed and dried under vacuum at 50°C for at least 12 hours. Vesicles were then formed at a lipid concentration of 5 mg/mL in a concentrated salt solution (1 mol/L NaCl, 100 mmol/L NaH_2_PO_4_, pH = 7.4). The solution was sonicated until it became translucent, typically after 90 minutes. The vesicles were allowed to incubate the tether SAM for 60 to 90 minutes before flushing with a low-salt buffer (100 mmol/L NaCl, 10 mmol/L NaH_2_PO_4_, pH = 7.4) to assist stBLM formation by osmotic shock. Rapid solvent exchange was used to form stBLMs for SPR (25). 50 µL of lipids in organic solvent (10 mg/mL) were added onto the tether SAM and incubated for ca. 1 min. The system was then flushed with at least 10 mL of the low salt buffer. For both procedures, the bilayers were flushed with subsequent rinses of 5 mL water, 10 mL ethanol/water (20 vol%), 5 mL water, and 10 mL of low salt buffer. The formation and quality of the stBLMs was assessed with electrochemical impedance spectroscopy (EIS) in terms of its bilayer resistance and capacitance (26), and SPR was used to quantify mass adsorption onto the membrane once the stBLMs were exposed to NTs in solution.

### Electrochemical Impedance Spectroscopy (EIS)

EIS measurements were performed between 1 Hz and 100 kHz, with 60 points logarithmically spaced, with a Solartron (Farnborough, UK) 1287A potentiostat and 1260 frequency analyzer in a three-electrode configuration. A saturated silver-silver chloride (Ag-AgCl-KCl(aq,sat)) microelectrode (M-401F; Microelectrode, Bedford, NH) served as the reference electrode with a 0.25 mm diameter platinum wire (99.9 % purity, Sigma Aldrich) coiled around the reference electrode as the auxiliary electrode. The working electrode was secured to the gold substrate with copper conducting tape (SPI Supplies/Structure Probe, Inc.; West Chester, PA). Data was collected and analyzed using Zplot and Zview (Scribner Associates, Southern Pines, NC) and fitted to models as described (27).

### Surface Plasmon Resonance

SPR measurements were performed at *T* = 25.00 ± 0.01°C in single-batch mode using two custom-built instruments that reflect light at wavelengths *λ* = 660 nm and 720 nm from the membrane-covered interfaces and work with glass and sapphire, respectively, as support media (SPR Biosystems, Germantown, MD). Gold-coated slides with a SAM layer were assembled by index matching to a prism (Kretschmann configuration), and stBLMs were prepared *in situ* as described above. The intensity of light reflected from the gold/buffer interface was recorded in a 2D-CCD detector as a function of time and the characteristic reflection minimum due to surface-plasmon generation, located on the detector at a position *R*, was determined in the raw signals using SPRAria (SPR Biosystems). To determine a baseline, the neat bilayer was measured for thirty minutes before adding NTs in increasing concentrations, *c*. *R*(*t*) was recorded for each concentration until equilibrium was approached at *R*_∞_ = *R*(*t*→∞) and the changes in *R*_∞_ as a function of *c* were fitted to a modified Langmuir isotherm:

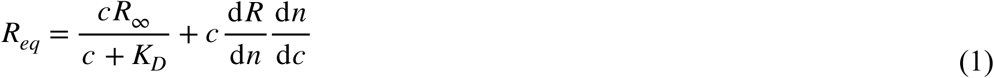

where *K*_*D*_ is the dissociation constant (28) and d*R*/d*n* is the instrument response to changes in refractive index of the bulk solution above the interface. The Langmuir binding model is based upon the assumption that all binding sites are equivalent, each ligand interacts with a single binding site, and ligands do not interact with each other. Because binding is expected to be weak, measurements were performed with millimolar NT concentrations which made corrections necessary, in the form of second term on the right hand side of Eq. (1), to account for changes in the refractive index of the solution. The refractive index increment, d*n*/d*c*, was determined using a prism refractometer at *λ* = 660 nm for blank buffer and known NT concentrations in solution.

For the quantitative interpretation of SPR signals, one may use the transfer matrix method (29) to calculate the shift of the optical reflection minimum in response to the adsorption of a molecular species to an interface whose structure is approximately known. The sample consists of a sequence of homogeneous layers, *i*, with thicknesses *d*_*i*_, refraction indices *n*_*i*_, and optical thickness *δ*_*i*_ = *d*_*i*_*k*_*i*_, where *k*_*i*_ is the wave vector component normal to the interface. The coefficients *r*_*n*,*n*+1_ and *t*_*n*,*n*+1_ then characterize reflection and transmission at each interface between subsequent layers and give rise to a transfer matrix element (30),

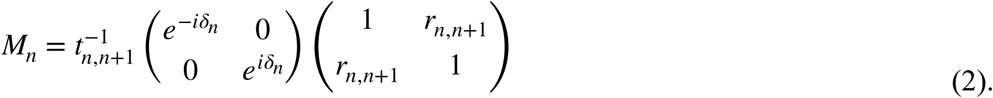

The product of the transfer matrices for all layers define the global optical properties of the composite film

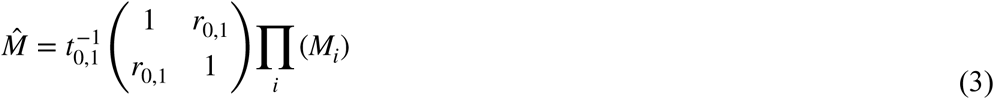

which relate its total reflectance and transmission as

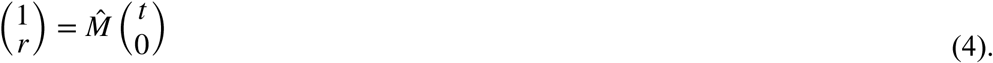

With an approximate structure of the interfacial film known from NR, and knowledge of the localization of adsorbed molecules with respect to bilayer components, which gives rise to estimates of the optical indices of the individual layers in the interfacial film, one can then relate shifts in SPR signal to mass adsorption, as shown in the results section.

### Neutron Reflectometry

NR is a sensitive method to characterize the organization of layered organic materials at interfaces and surfaces along their normal direction, *z* (31). Neutrons transmitted through the Si slab that supports an stBLM are reflected from its interface with the buffer at a shallow specular angle *α* with the momentum transfer *Q*_*z*_ = (4π/*λ*)⋅sin*α*, where the neutron wavelength *λ* is ≈ 5 Å. If the neutron scattering length densities (nSLDs) of substrate and adjacent liquid phase, *ρ*_substrate_ < *ρ*_buffer_, as is the case for Si bordering D_2_O-based buffer, the reflectivity *R* is unity between *Q*_*z*_ = 0 and 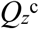, the critical momentum transfer for total internal reflection. Beyond 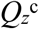, *R*(*Q*_*z*_) decays sharply, as almost all intensity is transmitted into the buffer. The reflection from an ideal interface follows the Fresnel reflectivity *R*_*F*_, which decays as 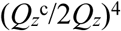 for sufficiently large *Q*_*z*_(32). If the idealized interface is blurred, usually modeled with a Gaussian roughness *σ*, the reflectivity decays even faster in *Q*_*z*_, as 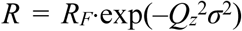. It is therefore essential to achieve ultra-low roughnesses of the substrate, such as *σ* = 3–5 Å on polished Si/SiO_2_.

A molecularly layered organic surface structure between the Si substrate and the buffer modifies *R*_*F*_ due to interferences between subsequent strata—the gold film, submembrane aqueous layer, membrane and surface-adsorbed molecules—and partial transmission and reflection amplitudes depend on the nSLDs, *ρ*, of individual layers. The neutron reflectivity can be approximated as 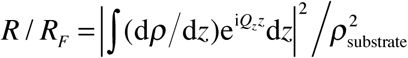(33). Due to the lack of phase information, the reflectivity cannot be directly inverted into a unique structural model. Yet, because the bilayer structure is approximately known, forward modeling of the data works well in practical terms.

Models of stratified interfacial structures of membranes on solid supports were traditionally constructed from molecular slabs of homogeneous nSLDs (box functions in the one-dimensional profiles, hence the frequently used term ‘box models’). More realistic descriptions are provided by models in which these strata are blurred into each other, parameterized as continuously variable distributions using analytic functions (34). Refinement of their parameters then approximates the underlying structure. Unique to the probing of interfacial structures with ‘cold’ neutron beams with kinetic energies in the meV range is the absence of beam damage, and thus the capability to perform subsequent measurements on the same sample. For example, as-prepared bilayers under neat buffer and bilayers exposed to NT in solution can be measured subsequently in isotopically distinct buffers. Thus, a sequence of individual measurements forms a large dataset, and their evaluation in context of each other restrains the parameters of the model strongly (35).

Measurements at *T* = 22 ± 1°C were performed on the NGD-Magik reflectometer (36) at the NIST Center for Neutron Research (NCNR). Reflectivity curves were recorded for momentum transfer values 0.001 ≤ *Q*_*z*_ ≤ 0.25 Å^−1^ for which counting statistics with adequate quality were typically obtained after 6 hours of beam-time per condition. A flow cell (volume ≈ 1.3 mL) allowed for *in situ* buffer exchange (37), accomplished by flushing the sample with ≈ 10 mL of buffer. Thereby, sequential measurements on membranes without and with NTs in H_2_O or D_2_O-based buffer were performed in repeated scans on the same sample footprint.

One-dimensional component volume occupancy (CVO) profiles (38) along the lipid bilayer normal (c.f. Fig. 1) were determined using a hybrid model that comprised a slab parameterization for the solid substrate (39), the continuous distribution of submolecular components for the stBLM (34), and a free-form model based on monotonic Hermite splines for adsorbed NTs. The model of the solid substrate was composed of slabs of bulk silicon, silicon oxide, chromium (deposited as a bonding layer), and the terminal gold film. Fit parameters were their thicknesses and nSLDs. For the stBLM, the submolecular groups were: interfacial βME, polyethylene glycol chains and glycerol groups of the HC18 SAM. HC18 alkyl chains were not distinguished from the acyl chains of the proximal phospholipid monolayer they intercalate. The phospholipid bilayers were parsed into their substrate-proximal and substrate-distal phosphatidylcholine and phosphatidylglycerol headgroups, substrate-proximal and substrate-distal polymethylene chains, and the chain-terminal methyls in the bilayer center. Many of these parameters are interdependent, and overall they were subject to the condition that they fill the available space, with water filling any volume not occupied by the surface architecture. Parameters fitted in the model were bilayer completeness, the surface-immobilized ratios of βME:HC18 and HC18:substrate-proximal phospholipid, the submembrane layer thickness, and the hydrocarbon layer thicknesses for each bilayer leaflet (Fig. 1). Volumes of the molecular components were constants in the model, and dependent parameters reveal system properties such as the area per phospholipid in the bilayer (38). Hermite splines, defined by control points separated by 15 Å on average, provided a model-independent description of the in-plane averaged NT distributions along *z*, interfacing seamlessly with the continuous distributions of stBLM components (34). Volume occupancies at the control points and deviations of their positions from equidistant separation were fitted parameters. A global roughness *σ* was applied to the substrate surface and all submolecular distributions. Parameter optimization was achieved using the *Refl1D* and *ga_refl* software packages developed at the NCNR (37). Fit parameters that were not expected to change across measurements, such as the structure of the solid substrate, were determined by a simultaneous fit of all reflectivity curves in a single dataset. Parameter confidence limits were determined by a Monte Carlo Markov Chain (MCMC)-based global optimizer (37).

## Results

Our investigations of NT interactions with phospholipid bilayers, reported below, pushed the limits of sensitivity of both the SPR characterization of adsorption thermodynamics and the structural characterization of NT/bilayer arrangements by NR. This required corrections for index changes of the bulk solution due to high NT concentrations in SPR measurements and a critical calibration of the NR model to account for deficiencies in the parametric descriptions of the interfacial structures. Both issues are described at the beginning of their respective sections.

### Surface Plasmon Resonance

To interpret SPR readings correctly at high concentrations of dissolved NT molecules which affect the optical index *n* of the buffer, their index increment, d*n*/d*c*, was measured. The neat buffer at *λ* = 660 nm had *n =* 1.3345. The addition of 100 mmol/L serotonin increased the index to *n* = 1.339 for d*n*/d*c* = 0.0021 ± 0.0001 dL/g, or (4.45 ± 0.21)×10^−5^ L/mmol. For GABA, d*n*/d*c* = 0.0017 ± 0.002 dL/g or (1.75 ± 0.21)×10^−5^ L/mmol.

Adsorption isotherms of two model NTs, serotonin and GABA, to stBLMs of different surface charges, in which POPC bilayers contained 0, 10, 30, or 50 mol% POPG, were determined with SPR. Representative data sets are shown in Fig. 2 and overall results listed in Table 1. All experiments were performed at least in triplicate and are quoted by their error-weighted means with uncertainties given as the error-weighted standard deviations. Serotonin adsorption occurred to all bilayer compositions and was detected at bulk concentrations above *c* ≈ 0.05 mmol/L. At *c* > 1 mmol/L, bulk phase index increases reached detectable levels, and a correction for d*n*/d*c* was necessary to fit the data. The corresponding models of the SPR data in Fig. 2A–C show the contributions of surface accumulation and bulk correction (red and blue lines in the on-line version of this article); the sum of the two (black lines) fit the experimental data well. For modeling of the entire isotherm, d*n*/d*c* was treated as a fit parameter, and ranged from (4.21 ± 0.16)×10^−5^ L/ mmol for POPC to (4.64 ± 0.21)×10^−5^ L/mmol for 7:3 POPC/POPG (Table 1), consistent with the value of d*n*/d*c* determined for serotonin in buffer by bulk index measurements. The affinity of serotonin to the bilayer surface is constant within experimental error near *K*_*D*_ = 1 mmol/L at low bilayer charge (0 and 10 mol% POPG) and high bilayer charge (50 mol% POPG). Remarkably, it drops by a factor of 2 for the bilayer that contains 30 mol% POPG (Fig. 3A), the concentration of charged phospholipids characteristic for the inner plasma membrane. The surface accumulation of serotonin, defined as *Γ*_∞_ = *R*_∞_(*c*→∞), was approximately constant within experimental error for all bilayer compositions (Fig. 3B).

**Table 1:** Fitted SPR results to quantify the adsorption serotonin, a membrane-affine NT, and GABA, a membraneinert NT, to stBLMs of different compositions. While the serotonin results required fitting to both arguments in Eq. (1), GABA results were well modeled by fitting only to the right hand side of Eq. (1).

| stBLM composition | PC | PC/PG 9:1 | PC/PG 7:3 | PC/PG 1:1 |
| --- | --- | --- | --- | --- |
| serotonin |  |  |  |  |
| dissociation constant, $K_D$ (mmol/L) | $1.1 \pm 0.2$ | $1.6 \pm 0.5$ | $0.42 \pm 0.16$ | $1.1 \pm 0.2$ |
| index change by surface accumulation, $R_\infty$ ($10^{-5}$) | $35 \pm 3$ | $30 \pm 5$ | $30 \pm 3$ | $35 \pm 3$ |
| bulk index change increment, $dn/dc$ (L/mmol) | $4.21 \pm 0.16$ | $4.45 \pm 0.15$ | $4.64 \pm 0.21$ | $4.29 \pm 0.12$ |
| model fit quality, $\chi^2$ | 2.122 | 1.549 | 2.351 | 0.966 |
| GABA |  |  |  |  |
| dissociation constant, $K_D$ (mmol/L) | n/a | n/a | n/a | n/a |
| index change by surface accumulation, $R_\infty$ ($10^{-5}$) | n/a | n/a | n/a | n/a |
| bulk index change increment, $dn/dc$ (L/mmol) | $1.77 \pm 0.12$ | $1.78 \pm 0.16$ | $1.66 \pm 0.10$ | $1.73 \pm 0.04$ |
| model fit quality, $\chi^2$ | 4.227 | 3.305 | 5.938 | 5.107 |

**Figure 2 (in color on-line):**
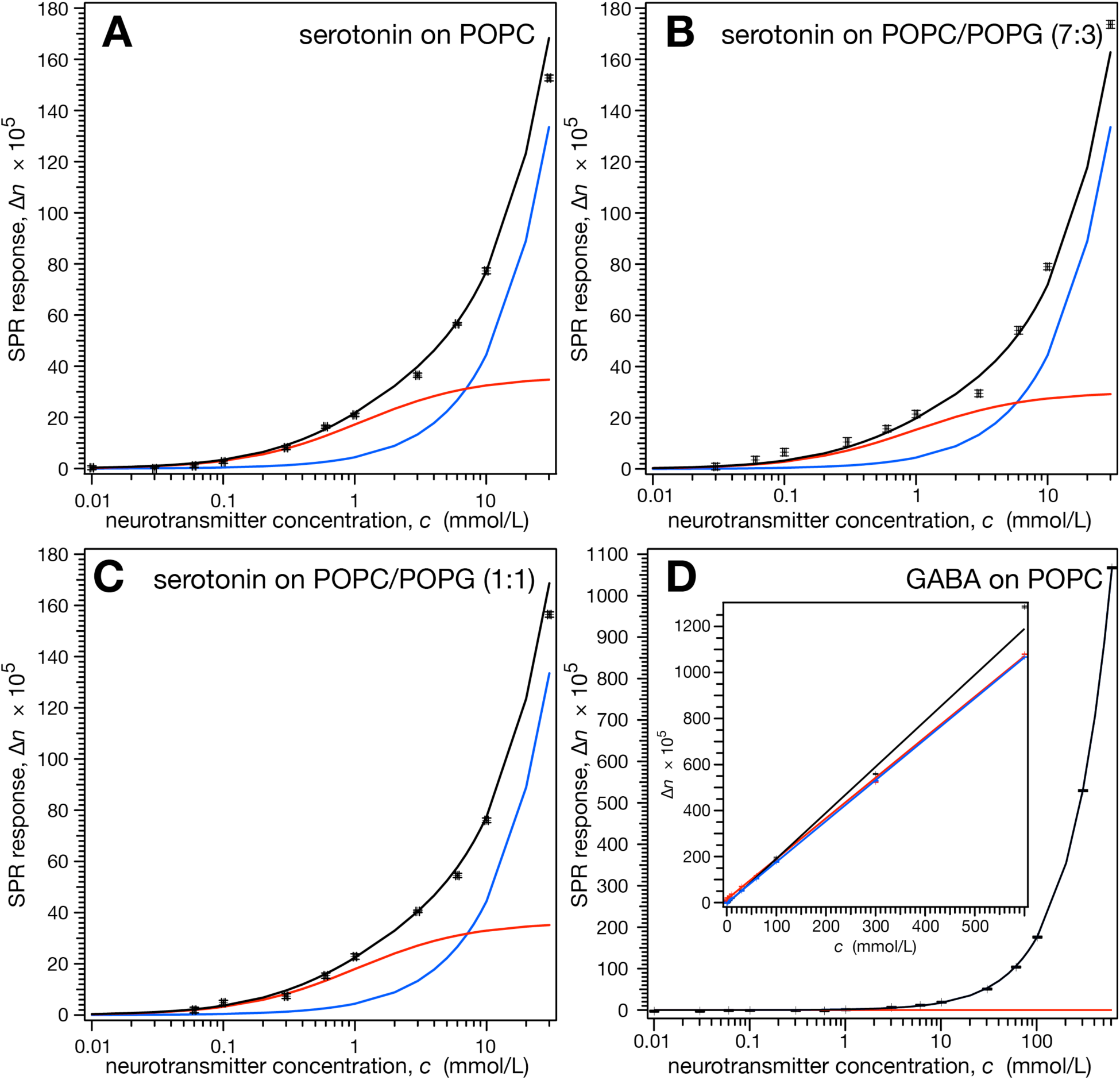
SPR measurements of stBLMs exposed to NTs in aqueous solutions. Equilibrium SPR responses, *R*(*t*➝∞), were converted into equivalent refractive index changes, Δ*n*, of the bulk solution adjacent to the membrane and are plotted in this form as a function of NT solution concentration. The overall signals (black) contain contributions from membrane adsorption and/or intercalation of NT molecules (red) and refractive index changes at high concentrations of dissolved NTs (blue). The two contributions are disentangled by fitting the data to Eq. (1). Panel D shows data for three independently measured bilayers on a linear scale.

**Figure 3:**
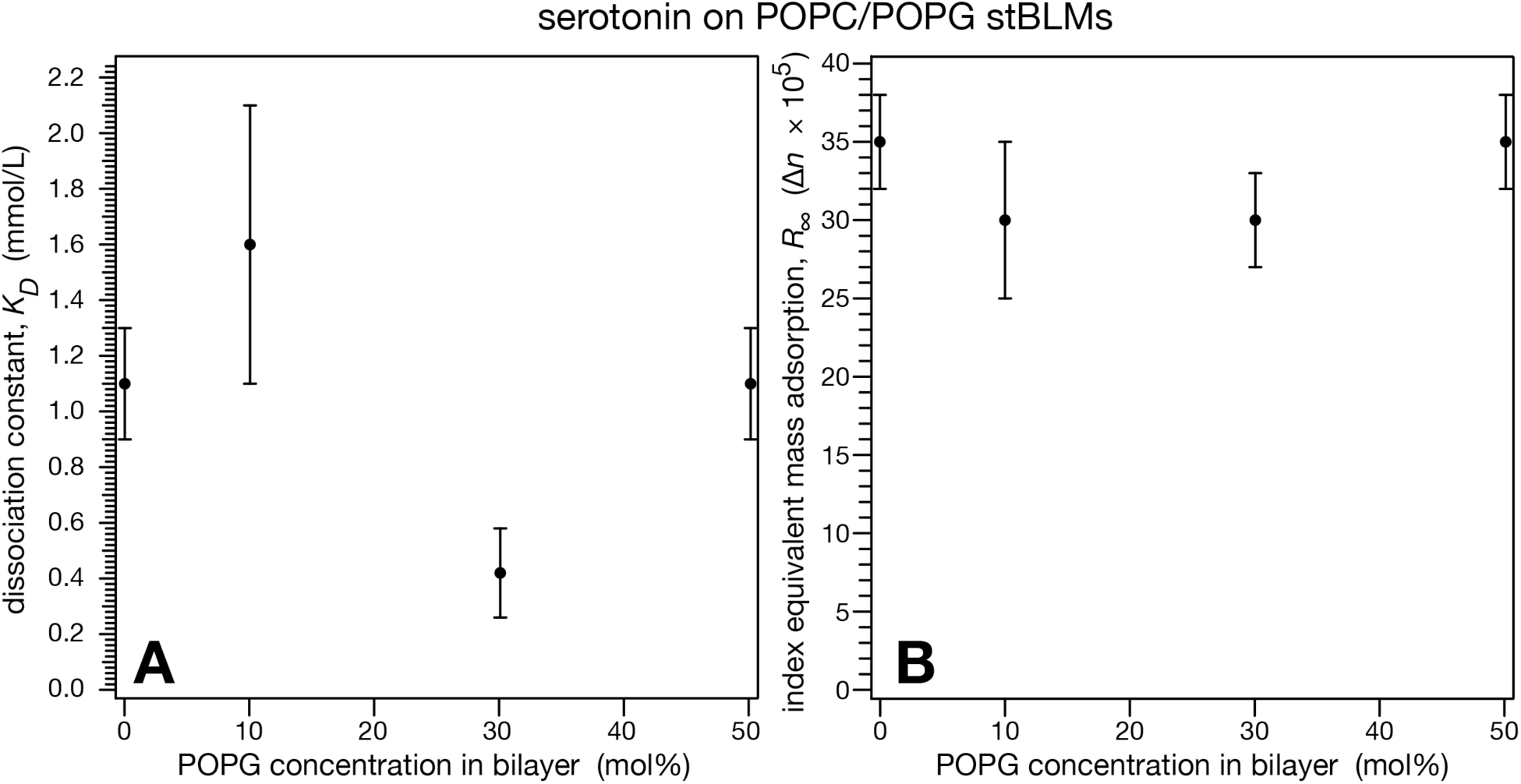
Comparison of serotonin binding to stBLMs that contain varying amounts of POPG. (A) The affinity (1/K_D_) shows a maximum at ≈ 30 mol% of the charged lipid, which coincides with the physiological concentration of anionic lipids—mostly phosphatidylserine—in the inner leaflet of the plasma membrane. (B) The extrapolated mass adsorption (*c*_NT_ ➝ ∞), displayed in equivalents of bulk index changes, is independent of bilayer surface charge.

In contrast to serotonin, the adsorption of the zwitterionic GABA to pure POPC or POPC/ POPG bilayers was below detection limits in our SPR experiments. Fits which only used the second term in Eq. (1)—*i.e.*, the bulk contribution to the signal—were entirely adequate to model the experimental data (Fig. 2D), with similar quality as the serotonin results fitted by the full adsorption model. This clearly implies that interfacial adsorption of GABA is below the sensitivity of the experiment.

### Neutron Reflectometry

NR was used to localize NTs adsorbed to stBLMs and quantify their adsorbed masses as a function of bilayer surface charge. Because the determination of adsorbed mass in SPR depends on the location of an adsorbed molecular species with respect to the bilayer, localization of adsorbed NT also permits *quantitative* estimates of the adsorbed mass as seen by the SPR experiments, and thus a cross-validation between the methods.

As in the SPR experiments, it was imperative to quantitatively assess the sensitivity of NR to NT adsorption to the interface in each measurement. To rationalize our sensitivity calibration procedure, we assume that there are systematic deviations from ideality of the sample structures in each individual measurement, such as a slight waviness of the substrate that will vary for each sample and is unaccounted for by data modeling. This implies that a description of the as-prepared stBLM—for which the model only contains the slab parameterization of the substrate and the continuous distribution of stBLM components, but no Hermite spline for an adsorbent—may be fraught with displacements of the best-fit parameters from their true values. If then a Hermite spline is added, to account for adsorption onto the stBLM in the modeling of the bilayer in the presence of NTs in solution, this free-form component will also populate to adjust for such systematic errors, *irrespective of whether or not molecules actually adsorb*. Differences in the structural models of pristine and NT-loaded stBLMs will thus contain contributions from molecular adsorption as well as contributions that may arise from model deficiencies. To assess the sensitivity of the experiment, we therefore determined the structure of the *as-prepared* stBLM with the model that allows for adsorbed molecules, assuming that the Hermite spline observed in this procedure represents a quantitative measure of the detection limits for an adsorbent on that particular sample.

We characterized serotonin association with stBLMs containing pure POPC and POPC/POPG (7:3) at various NT concentrations in the buffers. The concentration of the charged lipid component was chosen in view of the SPR results and the fact that charged phospholipids (mostly phosphatidylserine) account for ca. 30 mol% in the composition of the plasma membrane. Generally, the as-prepared bilayers were first measured in H_2_O-based and in D_2_O-based buffers to establish a baseline structure and calibrate the sensitivity of the ensuing experiments on bilayers under NT in solution. We then repeated the NR measurements on the same samples with serotonin at *c* = 1 and 10 mmol/L on POPC and *c* = 100 µmol/L to 10 mmol/L in three logarithmically spaced measurements for POPC/POPG. Fig. 4 shows an exemplary NR data set for 10 mmol/L serotonin. Differences between the as-prepared and NT-affected bilayer measurements are small but significant, as shown in the error-weighted residuals. Although counting errors are large at momentum transfers *Q*_*z*_ > 0.15 Å^−1^, the residual plots also demonstrate that there is information in the data up to ca. 0.25 Å^−1^.

**Figure 4 (in color on-line):**
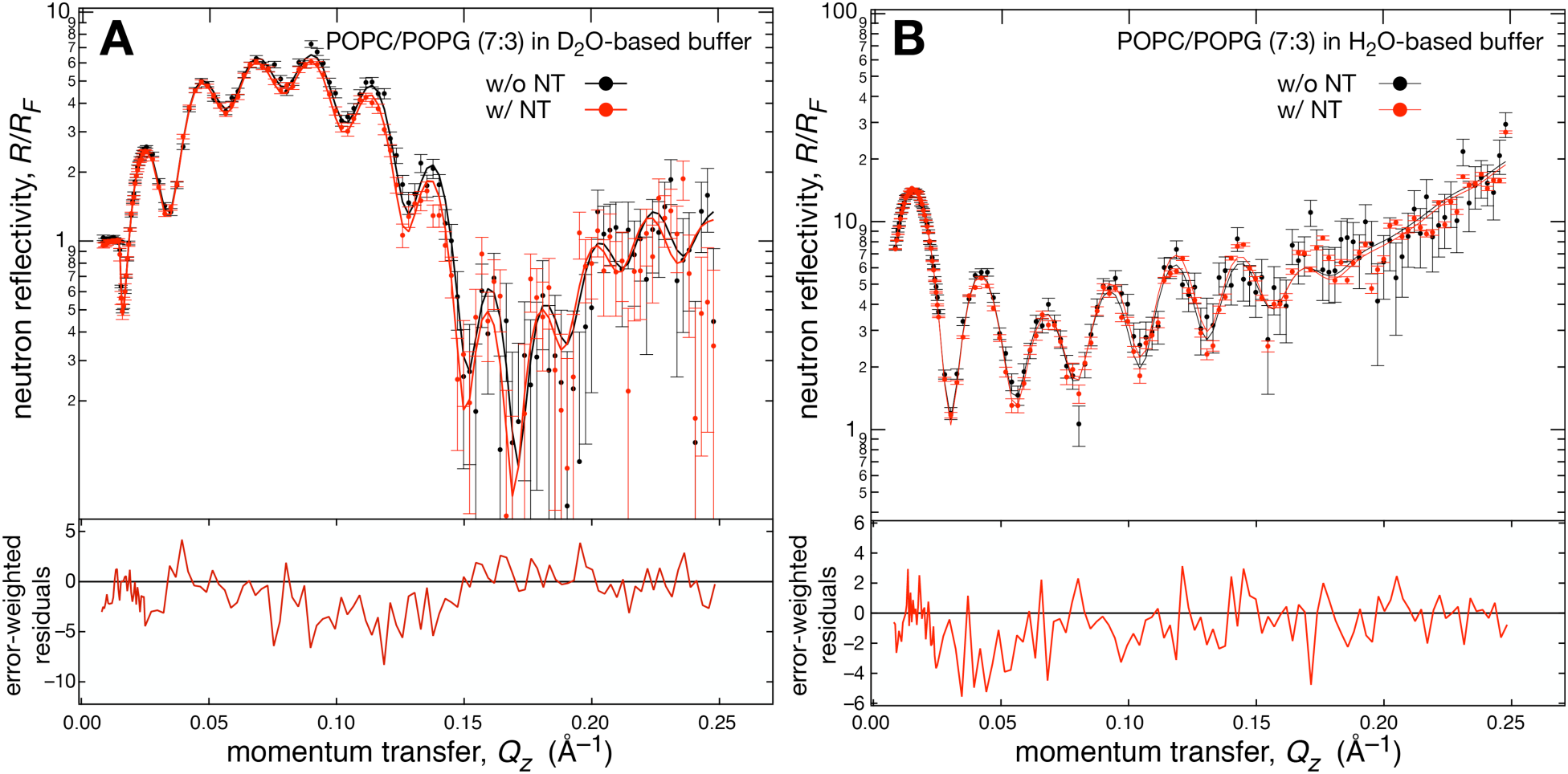
Neutron reflectivities of a 7:3 POPC/POPG stBLM before and after addition of 10 mmol/L serotonin in (A) D_2_O-based and (B) H_2_O-based buffer. Bottom: Error-bar weighted residuals show the differences between the NR curves measured with and without NT. All data sets were simultaneously evaluated and yielded a unique structural model, shown in Fig. 5D, in which adsorbed NT molecules were localized within the charged lipid bilayer.

Modeling of the data shown in Fig. 4 established a (combined) thickness of the hydrophobic chains, *d ≈* 29 Å, for the as-prepared bilayer, an average area per lipid, *A* ≈ 70 Å^2^, and a bilayer completeness of > 99% (first column in Table 2). Figure 5A shows an exemplary sensitivity calibration for a pure POPC bilayer which is typical for all samples studied. A blue line indicates adjustments of density resulting from model deficiencies; light blue areas show 68% confidence limits, and their upper bound provides a visual representation of the worst-case error. On this sample, this analysis established a resolution limit for serotonin of 6 ± 3 ng/cm^2^, above which we expect to be able to discern adsorbed NT.

**Table 2:** Model parameters from the structural characterization by NR of an as-prepared POPC stBLM and the same stBLM under buffer that contained serotonin. Median parameter values and 1*σ* (68%) confidence limits obtained from MCMC-based model refinement are shown. *The significance of a Hermite spline for the as-prepared bilayer lies in the fact that it determines deficiencies in the model that can be mistaken for adsorbed molecules on the same bilayer in contact with solutions containing NT. As such, this data provides an objective estimate of the sensitivity of the NR measurements to adsorbed molecular species (see Materials and Methods). ^†^Not significant

| parameter \ serotonin concentration | as prepared | 1 mmol/L | 10 mmol/L |
| --- | --- | --- | --- |
| bilayer completeness (%) | $99.5 \pm 0.5$ | $99.7 \pm 0.3$ | $99.6 \pm 0.4$ |
| area per lipid in distal leaflet, $A$ ($\text{\AA}^2$) | $69 \pm 3$ | $70 \pm 4$ | $67 \pm 3$ |
| total hydrocarbon thickness, $d$ ($\text{\AA}$) | $29.4 \pm 0.9$ | $29.4 \pm 0.9$ | $29.7 \pm 1.0$ |
| hydrocarbon thickness change, $\Delta d$ ($\text{\AA}$) | n/a | $0.0 \pm 1.3$ | $0.3 \pm 1.0$ |
| Hermite spline center of mass, $z$ ($\text{\AA}$) | $9 \pm 6^*$ | $10 \pm 7$ | $7 \pm 5$ |
| adsorbed NT mass per area ($\text{ng}/\text{cm}^2$) | $9.5 \pm 4.0^*$ | $9.6 \pm 4.0^\dagger$ | $15.2 \pm 5.6$ |
| NT/lipid in the substrate-distal leaflet | n/a | $0.23 \pm 0.10^\dagger$ | $0.35 \pm 0.13$ |
| model fit quality, $\chi^2$ | 1.17 | 1.06 | 1.10 |

**Figure 5:**
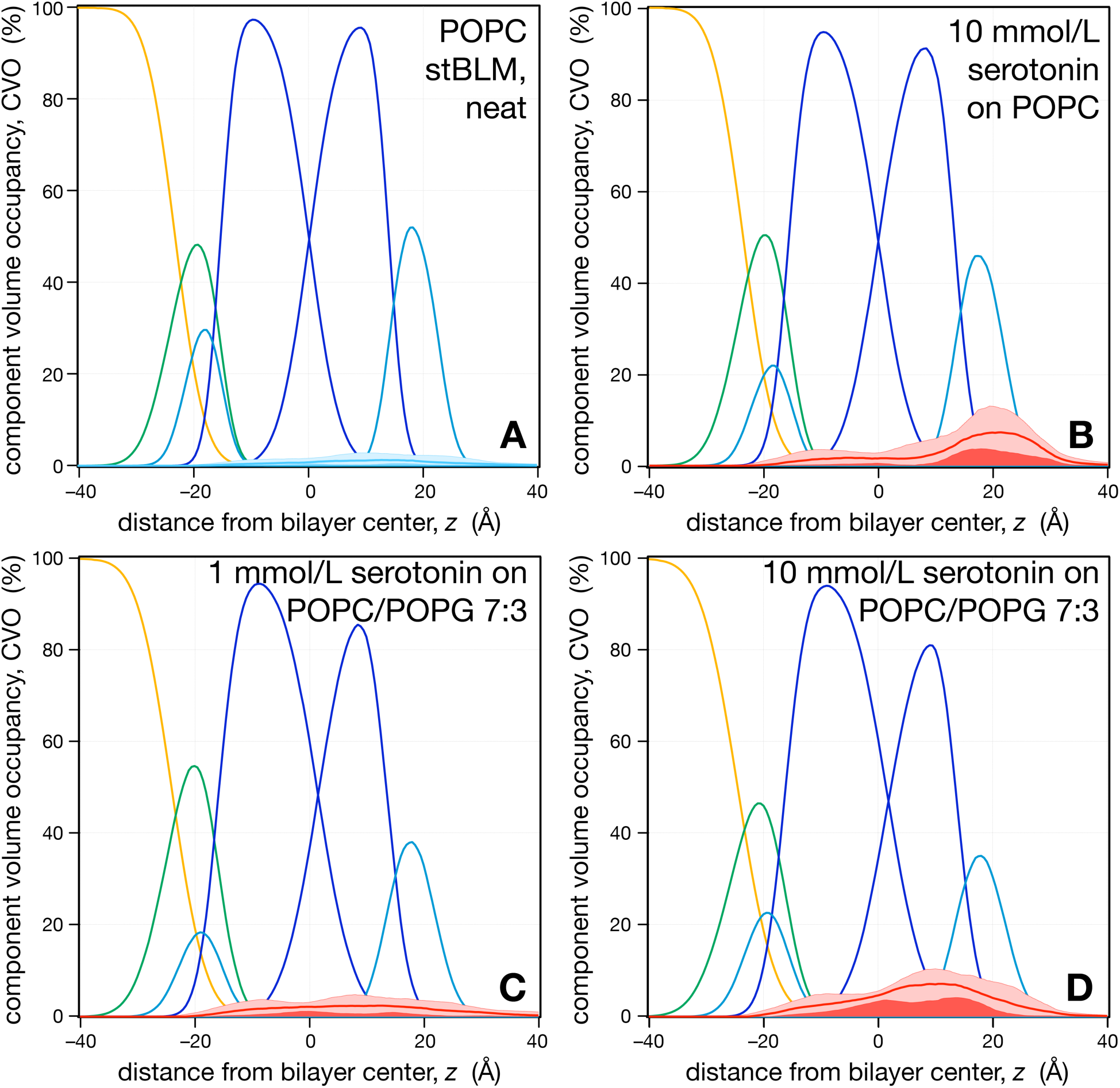
One-dimensional stBLM CVO profiles before and after incubation with serotonin. (A) Example of an as-prepared bilayer. The shaded blue area near the bottom of the panel visualizes model deficiencies that determine the sensitivity of the measurement to NT adsorption (see text). (B) A PC bilayer exposed to 10 mmol/L serotonin and (C, D) PC/PG (7:3) bilayers exposed to (C) 1 mmol/L and (D) 10 mmol/L serotonin. The color scheme is as in Fig. 1: Substrate-terminal gold layer: yellow; tether chemistry: green; lipid headgroups: turquoise; lipid chains: blue. Membrane-associated serotonin is shown in red with a line indicating the profile and shaded areas its 68% confidence limits. The solid red area visualizes minimal distributions of adsorbed NT when accounting for statistical and systematic errors.

Figure 5B–D shows representative CVO profiles of neutral and charged stBLMs (PC/PG = 7:3) in the presence of serotonin in the adjacent buffers. NT localized on the bilayers are shown in red, with a solid line for their distributions and light red areas indicating 68% confidence. Dark red areas visualize the minimum of likely distributions of NT molecules adsorbed to these bilayers in view of statistical and systematic errors, as described above. In particular, a comparison with Fig. 5A shows conclusively that the amounts of adsorbed NT exceed the sensitivity of the measurements. Figs. 5C and D show serotonin content in a charged bilayer upon incubation with serotonin at *c* = 1 and 10 mmol/L, respectively. These panels represent snapshots in a series of incubations with first 100 µmol/L, then 1 mmol/L, and finally 10 mmol/L, which resulted in serotonin mass adsorption between 1.8 ± 1.1 and 26 ± 5 ng/cm^2^ (Table 3). In response to this progression, we observed a ca. 10% decrease in the average area per lipid while the hydrophobic thickness remained constant. Bilayer completeness remained constant, suggesting that high concentrations of the NT do not induce defects in the membrane structure. A comprehensive list of the structural data determined for serotonin adsorption to stBLMs composed of POPC and POPC/POPG (7:3) is provided in Tables 2 and 3, respectively.

**Table 3:** Model parameters from the structural characterization by NR of an as-prepared stBLM composed of POPC/POPG (7:3) and the same stBLM under buffer that contained serotonin. Median parameter values and 1*σ* (68%) confidence limits obtained from MCMC-based model refinement are shown. *The significance of a Hermite spline for the as-prepared bilayer lies in the fact that it determines deficiencies in the model that can be mistaken for adsorbed molecules on the same bilayer in contact with solutions containing NT. As such, this data provides an objective estimate of the sensitivity of the NR measurements to adsorbed molecular species (see Materials and Methods). ^†^Not significant

| parameter \ serotonin concentration | as prepared | 0.1 mmol/L | 1 mmol/L | 10 mmol/L |
| --- | --- | --- | --- | --- |
| bilayer completeness (%) | 99.7 ± 0.3 | 99.7 ± 0.3 | 99.8 ± 0.2 | 99.8 ± 0.1 |
| area per lipid in distal leaflet, $A$ (Å<sup>2</sup>) | 83 ± 5 | 77 ± 3 | 77 ± 5 | 78 ± 6 |
| total hydrocarbon thickness, $d$ (Å) | 28.9 ± 0.9 | 27.8 ± 0.7 | 29.4 ± 1.1 | 29.8 ± 1.2 |
| hydrocarbon thickness change, $\Delta d$ (Å) | n/a | − 1.1 ± 1.3 | 0.5 ± 0.14 | 0.9 ± 1.6 |
| Hermite spline center of mass, $z$ (Å) | 16 ± 7* | 11 ± 8 | 9 ± 6 | 8 ± 3 |
| adsorbed NT mass per area (ng/cm<sup>2</sup>) | 5.7 ± 2.4* | 1.8 ± 1.1<sup>†</sup> | 13.0 ± 3.9 | 26.0 ± 5.1 |
| NT/lipid in the substrate-distal leaflet | n/a | 0.05 ± 0.03<sup>†</sup> | 0.34 ± 0.11 | 0.70 ± 0.14 |
| model fit quality, $\chi^2$ | 1.47 | 1.38 | 1.39 | 1.59 |

Intriguingly, serotonin was primarily localized within the bilayer upon adsorption to the membranes, essentially coinciding with the distribution of the substrate-distal phospholipids. Already at bulk serotonin concentrations *c <* 1 mmol/L, NT molecules were detected close to the interface between lipid hydrocarbon tails and headgroups of the distal monolayer. Upon incubation with subsequently higher concentrations, the amount of the trapped NT increased, primarily in the outer leaflet (Fig. 5C,D). For the exposure of pure POPC stBLMs to serotonin at high concentration (*c* = 10 mmol/L), NR showed a similar distribution of the serotonin CVO profiles, but the amount of adsorbed material was slightly lower than in the charged bilayers (Fig. 5B and Table 2).

We also measured the structures of stBLMs composed of pure POPC and POPC/POPG (7:3) at various concentrations of GABA up to 100 mmol/L in the adjacent buffer. Sensitivity calibrations of the corresponding real-space models established a sensitivity limit of 10 ng/cm^2^ for GABA (Fig. 6A and Table 4). In all models for bilayers incubated with GABA, we observed minute changes of the structural parameters, but the amounts of adsorbed or incorporated neurotransmitter were at best borderline significant, as shown for 10 mmol/L of GABA on a zwitterionic and a charged bilayer in Figs. 6B and C, respectively. A comprehensive list of the structural data determined for GABA adsorption to stBLMs composed of POPC and POPC/POPG (7:3) is provided in Tables 4 and 5, respectively.

**Table 4:** Model parameters from the structural characterization by NR of an as-prepared POPC stBLM and the same stBLM under buffer that contained GABA. Median parameter values and 1*σ* (68%) confidence limits obtained from MCMC-based model refinement are shown. *The significance of a Hermite spline for the as-prepared bilayer lies in the fact that it determines deficiencies in the model that can be mistaken for adsorbed molecules on the same bilayer in contact with solutions containing NT. As such, this data provides an objective estimate of the sensitivity of the NR measurements to adsorbed molecular species (see Materials and Methods). ^†^Not significant

| parameter \ GABA concentration | as prepared | 10 mmol/L | 100 mmol/L |
| --- | --- | --- | --- |
| bilayer completeness (%) | 99.7 ± 0.3 | 99.7 ± 0.3 | 99.7 ± 0.3 |
| area per lipid in distal leaflet, $A$ (Å<sup>2</sup>) | 68 ± 3 | 72 ± 4 | 70 ± 3 |
| total hydrocarbon thickness, $d$ (Å) | 29.9 ± 0.9 | 29.9 ± 1.0 | 29.4 ± 0.9 |
| hydrocarbon thickness change, $\Delta d$ (Å) | n/a | 0.0 ± 1.0 | − 0.5 ± 1.3 |
| Hermite spline center of mass, $z$ (Å) | 10 ± 4* | 10 ± 5 | 10 ± 6 |
| adsorbed NT mass per area (ng/cm<sup>2</sup>) | 9.7 ± 4.1* | 11.5 ± 6.2<sup>†</sup> | 9.9 ± 4.8<sup>†</sup> |
| NT/lipid in the substrate-distal leaflet | n/a | 0.49 ± 0.26<sup>†</sup> | 0.41 ± 0.20<sup>†</sup> |
| model fit quality, $\chi^2$ | 1.15 | 1.01 | 0.98 |

**Table 5:**
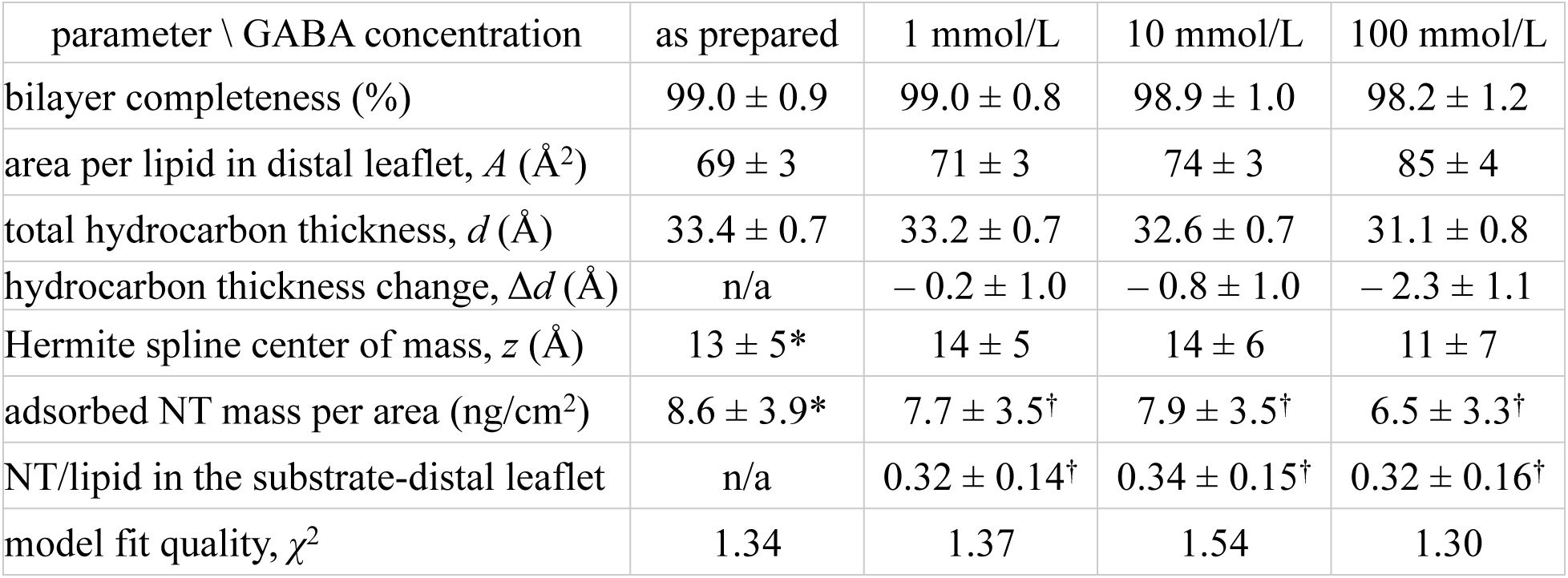
Model parameters from the structural characterization by NR of an as-prepared stBLM composed of POPC/POPG (7:3) and the same stBLM under buffer that contained GABA. Median parameter values and 1*σ* (68%) confidence limits obtained from MCMC-based model refinement are shown. *The significance of a Hermite spline for the as-prepared bilayer lies in the fact that it determines deficiencies in the model that can be mistaken for adsorbed molecules on the same bilayer in contact with solutions containing NT. As such, this data provides an objective estimate of the sensitivity of the NR measurements to adsorbed molecular species (see Materials and Methods). ^†^Not significant

**Figure 6:**
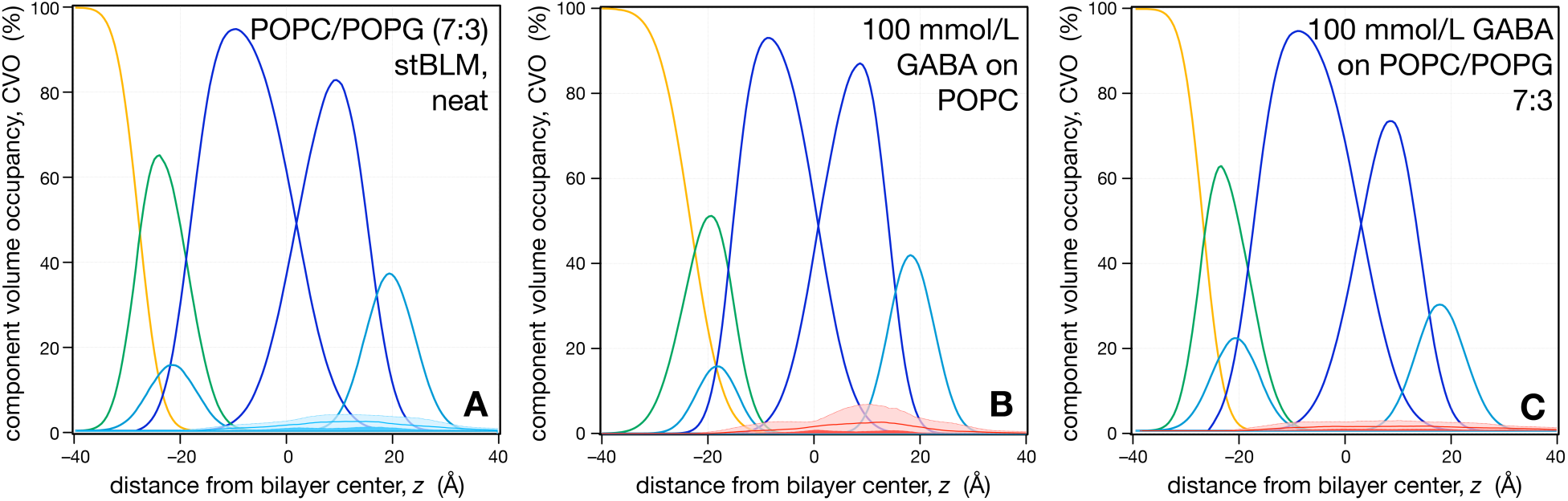
One-dimensional stBLM CVO profiles before and after incubation with GABA. (A) Example of an asprepared bilayer. (B, C) A PC bilayer (A) and a PC/PG (7:3) bilayer (C) exposed to 100 mmol/L GABA. Details as in Fig. 5.

### Quantitative evaluation of SPR data

SPR and NR both provide quantitative accounts of adsorbed NT on stBLM bilayers, as shown in Figs. 2 and 5-6. However, while the NR quantification of adsorbed NT can be directly derived from integration of the CVO components, a quantitative interpretation of the SPR results depends on the position of the adsorbed molecules with respect to the bilayer, because the same NT mass superficially associated with the bilayer results in a larger contribution to the SPR signal than the same NT mass intercalated into the hydrophobic core. Because this structural information is not inherent to the SPR data, we were not able to quantify NT mass adsorption (Fig. 2) without reference to an independent structural method. With the structural information obtained from NR (previous section), we can now revisit this issue and determine an absolute measure of the adsorbed molecular mass as seen by SPR.

The shift in the minimum of the SPR curve, Δ*θ*_*SPR*_, was calculated using the matrix transfer mechanism (Eqns. 2–4) for a model system composed of 5 layers between the Si wafer and bulk water that represented the substrate and stBLM. The substrate was composed of either a glass or sapphire base for the *λ* = 660 nm and 720 nm instruments, respectively, topped with chromium and gold layers. A simplified stBLM model included a water-rich tether, substrate-proximal and substrate-distal lipid monolayers, each assumed to be homogeneous within, and adjacent buffer. The thickness of individual layers matched the sputtering protocol and NR fits, and optical properties were determined from reference tables (40–43). While the refractive indices and dispersion coefficients of the substrate differed for the two instruments, they were constant for the stBLMs within the validity of our assumptions. To estimate the refractive index of a serotonin-intercalated monolayer, the index of the lipid layer (*n* = 1.45, ref. (44)) was modified by the addition of serotonin with an index estimated at *n* = 1.491 from the Lorentz-Lorenz relation

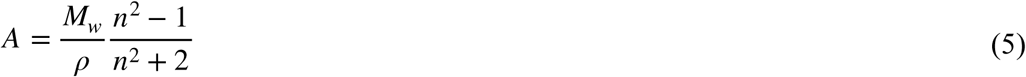

where *A* = 51.021 is the molar refractivity determined from the chemical structure (45), and *M*_*w*_ and *ρ* are the NT molar mass and density, respectively. While the validity of Eq. (5) is limited to non-interacting media, this condition is satisfied because NT affinities to the bilayer are low. Our analysis led to a volume-weighted value of *n* = 1.4595 if one assumes a mass adsorption of ≈ 37 ng/cm^2^ to the surface, equivalent to 2 NT molecules per lipid in the substrate-distal leaflet. The proximal leaflet was kept at *n* = 1.4500, in accord with our observation with NR that serotonin only intercalates the substrate-distal bilayer leaflet. Using this interpretation of the SPR data to determine the amount of NT adsorbed as a function of bulk concentration shows that both SPR and NR results are consistent and follow a Langmuir isotherm, see Fig. 7. In distinction, masses of serotonin consistent with NR but adsorbed on the membrane surface in a hydrated layer would give rise to SPR signals by a factor of 4 greater than those shown. This alternate model therefore underestimates the adsorbed NT mass significantly and is inconsistent with the NR results.

## Discussion

We measured the non-specific adsorption of two NTs to bilayers composed of POPC and POPG and observed that serotonin associates with the membranes to a significant extent while GABA adsorption is below our sensitivity limits. SPR measurements on planar bilayers to establish *K*_*D*_ and *R*_∞_ are appropriate due to the high sensitivity of the devices, ≈ 3×10^−7^ by refractive index (46). This sensitivity is achieved by using atomistically flat substrates with RMS roughnesses, σ < 5 Å. As demonstrated in Fig. 2, fitting neurotransmitter adsorption to Eq. (1) models the data accurately by separating an increase in SPR response due to bilayer adsorption from that due to changes in the bulk index (47). In the case of serotonin adsorption to PC and PC/PG bilayers, the bulk effect is small up to *c* ≈ 1 mmol/L, where *R*_∞_ ≈ 0.5 ng/cm^2^. There is a crossover of bulk and bilayer effects at *c* < 10 mmol/L after which the bulk effect dominates surface adsorption. Serotonin surface accumulation was verified with NR, which is inherently surface-sensitive (32). Beyond quantification of surface accumulation, however, NR provides further information on the localization of the neurotransmitters within the bilayers.

**Figure 7 (in color on-line):**
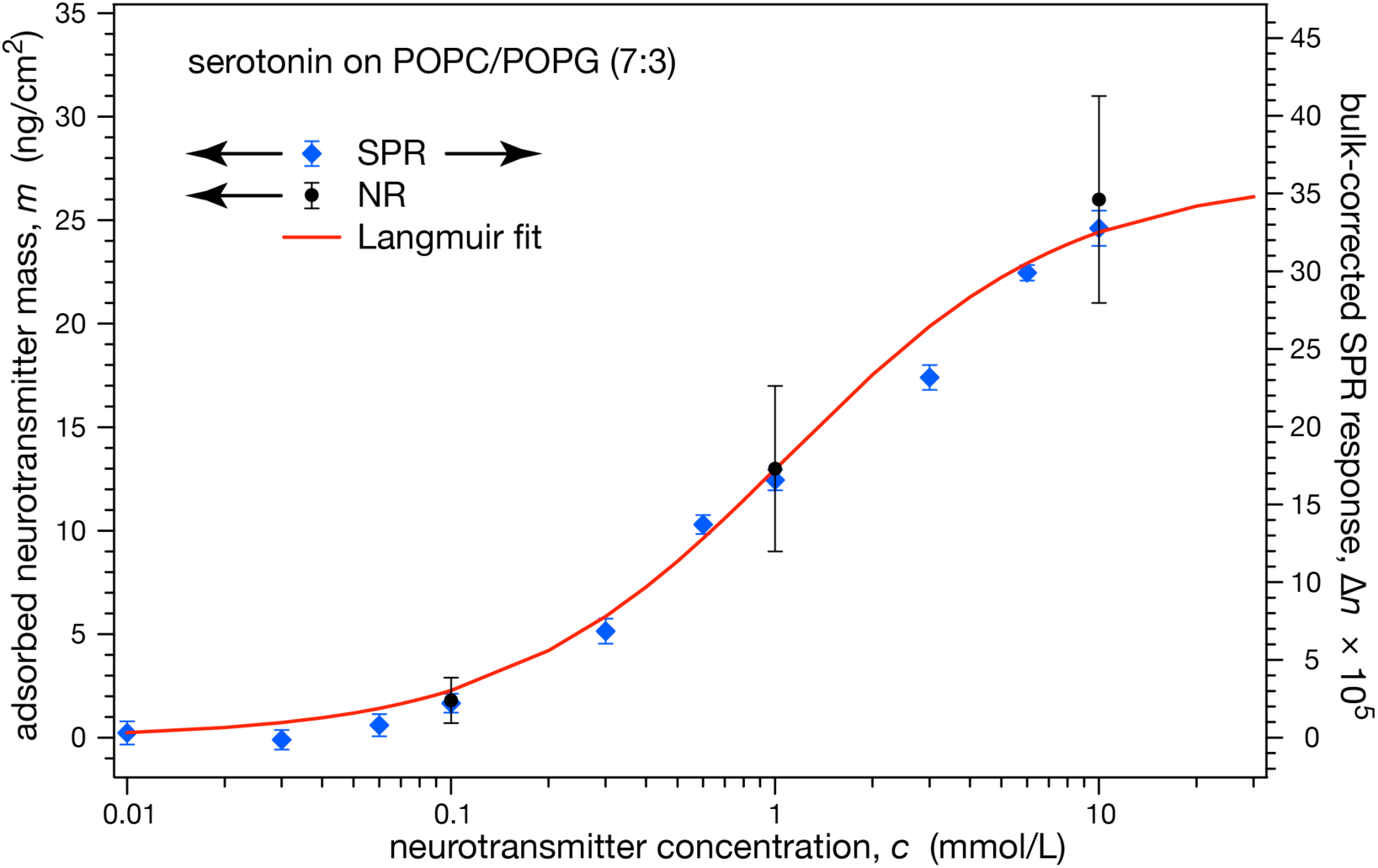
Comparison of adsorbed NT mass (left scale) as determined from SPR (blue diamonds) and NR (black dots) measurements of a PC/PG (7:3) stBLMs exposed to 10 mmol/L serotonin in aqueous buffer. The best fit Langmuir adsorption isotherm, Eq. (1) without the bulk contribution for dissolved NT, is shown in red for comparison (right scale).

In line with predictions from MD simulation (19), the binding characteristics of the NTs chosen for this study differ significantly. GABA does not show measurable affinities to bilayers composed of PC or PC/PG while serotonin interacts with such bilayers to a significant extent, and embeds deeply. Electrostatic interaction contributes to the recruitment of serotonin to charged bilayers, as we observe a drop of *K*_*D*_ near 30 mol% PG, although its surface load, *R*_∞_, remains approximately constant between 0 and 50 mol% of PG (Fig. 3). This confirms a tendency observed in MD simulations (19), where the addition of 20% PS to POPC in a bilayer also reduced *K*_*D*_ by about 50%, as estimated from the free energy profiles reported there, but in absolute terms, our measured *K*_*D*_ values are lower by a factor of ≈ 10 than those inferred from ref. (19). Distinct from serotonin, the SPR response for GABA was linear over a broad concentration range, between 0 and 1 mol/L, with a slope consistent with the concentration-dependent increase of the bulk index. Indeed, fitting of the GABA adsorption isotherms to the full Eq. (1) did not yield meaningful results, as the parameter uncertainties were typically larger than the parameter values. In essence, the total amount of GABA adsorbed to the bilayer was below the instrumental resolution, estimated from the uncertainties in *R*_∞_ at 5×10^−2^ ng/cm^2^.

Our NR results show serotonin adsorbed to the hydrophilic-hydrophobic interface between the lipid headgroup and hydrocarbon tails of the exposed leaflet of a PC bilayer. This, again, agrees with MD simulations which indicated that serotonin may be hydrogen-bonded to lipid phosphates and observed the long NT axis oriented perpendicular to the bilayer (14). A careful analysis of our NR-based attempts to quantify GABA association with phospholipid bilayers showed that any changes in the membrane structure that might be attributed to NT accumulation is below, or at most equal to, systematic uncertainty, estimated at 10 ± 4 ng/cm^2^ (Tables 4 and 5). This is consistent with the SPR results as well as with earlier equilibrium binding experiments that evaluated GABA binding to DMPC liposomes by dialysis (13). However, our measurements do not confirm other findings in that paper, as we did not detect any GABA association with anionic bilayers. On the other hand, our GABA results are consistent with MD simulations which observed that this NT remains excluded from both neutral and charged bilayers (19). The data reported here put an lower limit on GABA membrane affinity that corresponds to *K*_*D*_ ≈ 300 mmol/ L. This exceeds by far GABA’s peak concentration, ≈ 5 mmol/L, in the synaptic cleft (48, 49).

The differences in bilayer affinities of serotonin and GABA correlate with their distinct chemical characteristics and can also be rationalized by the distinct localizations of the binding sites on their postsynaptic receptor proteins. Serotonin is hydrophobic, so insertion into the bilayer is more favorable than for the more hydrophilic GABA. This hydrophobicity effect has been observed for other small molecules that partition into the bilayer, for example in the form of the Meyer-Overton rule which relates the partitioning of volatile anesthetics in mineral oil to their physiological efficacy (50, 51). The hydrophobic NT melatonin is structurally related to serotonin and has been reported to adsorb to bilayers and intercalate at a similar depth (16).

Our results on GABA suggest that it does not partition to bilayers at concentrations assumed to be relevant by Lee *et. al* (9) in their modeling of electrophysiology data. In line with our findings, the ligand binding site on the GABA_A_ receptor is exposed to the synaptic cleft (52). Therefore, GABA adsorption to the bilayer, even if a weak effect, would compete with its binding to the receptor. In contrast, serotonin receptor proteins have binding sites buried in the membrane (53). Thus non-specific membrane adsorption that enables 2D diffusion once a NT hits the surface would decrease the overall diffusion time to its binding site in comparison to a purely 3D random walk in the aqueous environment (54). In addition to this effect of membrane adsorption on binding kinetics to its cognate receptor, the unspecific binding of serotonin to the bilayer can elicit indirect affects on transmembrane proteins as they might alter the lateral pressure profile across the membrane (11, 55). An indication that this may indeed be relevant is our observation that neither membrane thicknesses nor areas per lipid increase proportionally to the NT volumes inserted into the bilayer. Serotonin has a molecular volume which is about 20% of that of phospholipids (≈ 240 Å^3^ vs. ≈ 1150 Å^3^) and its adsorbed mass saturates at ca. 0.7 NT molecules per membrane lipid (Table 3). Yet we observe no significant volume changes of the membrane, suggesting that the inserted volume can only be accommodated by lateral pressure increases, which due to the inhomogeneous distribution of intercalated material across the membrane would in turn suggest changes in the pressure profile. While our observations make such changes likely, of the two NT molecules studied here this only applies to serotonin but probably not for GABA. Nevertheless, theoretical (8, 11, 56) and computational (57) approaches and, often indirectly, experimental evidence (58–61) suggest that changes in the pressure profile within the bilayer can affect the functionality of transmembrane proteins, as well as the association, and thus the functionality, of peripheral membrane proteins (55, 62). Our results from the work reported here demonstrate conclusively that serotonin binds to zwitterionic and charged membranes with significant affinities. The *K*_*D*_ values we determined that are by far lower than serotonin concentrations in the intercellular space after ejection from the presynaptic dendrite. In extrapolation to a class of NTs with moderate hydrophobicity, characterized as membrane-affine in comprehensive MD simulations (19), it is thereby likely that their association with the post-synaptic membrane affect the dynamic response of membrane receptors by mechanisms other than the specific binding to their cognate receptors.

## Conclusions

We measured the non-specific adsorption of two neurotransmitters to model membranes and found that serotonin adsorbs at physiological concentrations while GABA does not. SPR experiments determined membrane dissociation constants, *K*_*D*_, in the millimolar range for serotonin, matching its concentrations upon ejection into the synaptic cleft. In contrast, GABA was not observed to adsorb to neutral or charged membranes up to bulk concentrations approaching 1000 mmol/L. NR experiments confirmed these findings and showed that serotonin intercalates into the distal leaflet of the bilayer where it is centered near the lipid headgroup phosphates. GABA adsorption could not be quantified in these measurements above the intrinsic detection limits. The two experimental methods were cross-validated by simulating the SPR response of a bilayer with the structural results obtained from NR. Indeed, the results of the two techniques were consistent with each other only if we assumed that serotonin intercalates the bilayer in an optical model that interpreted the SPR results quantitatively. This evidence suggests that aromatic neurotransmitters adsorb to the postsynaptic membrane, as predicted by atomistic MD simulations (19). Their presence in the bilayer is likely to alter the energetic landscape of the membrane-embedded receptor proteins and should be taken into account when modeling electrogenic responses of the postsynaptic membrane (11).

## Author contributions

F.H. and M.L. designed the research, B.J. and F.H. prepared samples and performed experiments, B.J., F.H., V.S. and M.L. analyzed data, B.J., F.H. and M.L. wrote the manuscript and all authors approved the final version.

## Acknowledgments

We thank David J. Vanderah for HC18 and Frederick Lanni for advice on the quantitative interpretation of SPR results. Neutron beamtime obtained at the NIST Center for Neutron Research is gratefully acknowledged. This work was supported by the U.S. Department of Commerce through grant no. 70NANB17H299.

Certain commercial materials, equipment, and instruments are identified in this manuscript in order to specify the experimental procedure as completely as possible. In no case does such identification imply a recommendation or endorsement by the National Institute of Standards and Technology, nor does it imply that the materials, equipment, or instruments identified are necessarily the best available for the purpose.

## Note

The authors declare no conflicts of interest.

